# Brewpitopes: a pipeline to refine B-cell epitope predictions during public health emergencies

**DOI:** 10.1101/2022.11.28.518301

**Authors:** Roc Farriol-Duran, Ruben López-Aladid, Eduard Porta-Pardo, Antoni Torres, Laia Fernández-Barat

## Abstract

The application of B-cell epitope identification for the development of therapeutic antibodies is well established but consuming in terms of time and resources. For this reason, in the last few years, the immunoinformatic community has developed several computational predictive tools.

While relatively successful, most of these tools only use a few properties of the candidate region to determine their likelihood of being a true B-cell epitope. However, this likelihood is influenced by a wide variety of protein features, including the presence of glycosylated residues in the neighbourhood of the candidate epitope, the subcellular location of the protein region or the three-dimensional information about their surface accessibility in the parental protein.

In this study we created Brewpitopes, an integrative pipeline to curate computational predictions of B-cell epitopes by accounting for all the aforementioned features. To this end, we implemented a set of rational filters to mimic the conditions for the *in vivo* antibody recognition to enrich the B-cell epitope predictions in actionable candidates. To validate Brewpitopes, we analyzed the SARS-CoV-2 proteome. In the S protein, Brewpitopes enriched the initial predictions in 5-fold on epitopes with neutralizing potential (p-value < 2e-4). Other than S protein, 4 out of 16 proteins in the proteome contain curated B-cell epitopes and hence, have also potential interest for viral neutralization, since mutational escape mainly affects the S protein. Our results demonstrate that Brewpitopes is a powerful pipeline for the rapid prediction of refined B-cell epitopes during public health emergencies.

**Statement of significance:** We have created Brewpitopes, a new pipeline that integrates additional important features such as glycosylation or structural accessibility, to curate B-cell epitope more likely to be functional in vivo. We have also validated Brewpitopes against SARS-CoV-2 not only for S protein but also for the entire viral proteome demonstrating that is a rapid and reliable epitope predictive tool to be implemented in present or future public health emergencies. Brewpitopes has identified 7 SARS-CoV-2 epitopes in S and epitopes allocated in 4 other proteins. Overall, offering an accurate selection of epitopes that might be scaled up to the production of new antibodies.

## Introduction

Neutralizing antibodies play a major role in the adaptive immune response against pathogens^1^. Hence, the prediction of the protein regions these recognize is key to guide the understanding of their mechanism of action ^1^. These protein regions, termed B-cell epitopes, potentially spread through the entire proteome of the target organism. This wide distribution requires of high throughput techniques to unravel the full epitope landscape. In this context, the bioinformatic prediction of B-cell epitopes has stepped in as a necessary exploration prior to the experimental validation (Table 1). For instance, in the race against SARS-CoV-2 pandemics, accurate bioinformatic B-cell epitope predictors significantly contributed to the success of COVID-19 preventive and therapeutic strategies ^2^ (Table 2). For this reason, many groups dedicated efforts to the identification of SARS-CoV-2 antibody binding regions using different bioinformatic approaches as a first step to later characterize neutralizing antibodies or to design immunogens for vaccines (Table 2) ^2,3^.

**Table 1.** Comparison of the state-of-the-art B-cell epitope predictors. Here, we have classified the state-of-the-art B-cell epitope predictions according to the features included such as subcellular location, glycosylation, or surface accessibility. Additionally, we also noted the dataset they are trained on.

| Method | Type of Epitope | Subcellular location | Glycosylation | Surface Accessibility | Based on | Comment | Reference |
| --- | --- | --- | --- | --- | --- | --- | --- |
| Bepipred 2.0 | Linear | No | No | Yes | Ab-Antigen structures | Buried residues are predicted within epitopes. | 5 |
| ABCpred | Linear | No | No | No | BCIPEP | High numbers of epitopes due to different window lengths. | 16, 29 |
| EpitopeVec | Linear | No | No | No | IEDB + BCIPEP | - | 7, 20, 29 |
| SVMTriP | Linear | No | No | No | IEDB | - | 8, 20 |
| Discotope 2.0 | Conformational | No | No | Yes | Ab-Antigen structures | - | 6, 20 |
| SeRenDIP-CE | Conformational | No | No | No | SAbDab | - | 9, 12 |
| SEPPA 3.0 | Conformational | Yes | Yes | No | Ab-Antigen structures | Predefined subcellular location. Does not take into account protein regions and does not include O-glycosylations. | 10 |
| Ellipro | Conformational | No | No | Yes | Antigen structures | - | 11 |
| Epitope3D | Conformational | No | No | No | Ab-Antigen structures | - | 13 |
| Brewpitopes | Linear + Conformational | Yes | Yes | Yes | Ab-Antigen structures | - | - |

**Table 2.** Comparison of the B-cell epitope studies of SARS-CoV-2. Here, we have collected the different B-cell epitope prediction approaches that have studies the proteome of SARS-CoV-2. The different publications were classified according to the features included such as subcellular location, glycosylation, or surface accessibility. Additionally, we also noted the B-cell epitope predictors and the techniques they implemented.

| Authors | Type of Epitope | Epitope Predictor | Subcellular location | Glycosylation | Surface Accessibility | Based on | Comment | Reference |
| --- | --- | --- | --- | --- | --- | --- | --- | --- |
| Almofti et al. | Linear + Conformational | IEDB | No | No | No | IEDB | - | 37 |
| Ezaj et al. | Linear + Conformational | IEDB | No | No | No | IEDB | - | 29 |
| Sikora et al. | Conformational | - | No | Yes | No | Molecular dynamics | - | 30 |
| Khare et al. | Conformational | - | No | No | Yes | Homology modelling | - | 31 |
| Smith et al. | Linear | - | No | Yes | Yes | - | Integrates sequence conservation and functional domains. | 32 |
| Li et al. | Linear | - | No | No | No | Epitope mapping |  | 33 |
| Schwarz et al. | Linear | - | No | No | No | Epitope mapping |  | 34 |
| Grifoni et al. | Linear | IEDB | No | No | Yes | Homology modelling | - | 35 |
| Haynes et al. | Conformational | SERA | No | No | No | Bacterial display libraries | - | 36 |
| Farriol-Duran et al. | Linear + Conformational | Multiple | Yes | Yes | Yes | Ab-Antigen structures | - | - |

B-cell epitope predictors recommended by the Immune Epitope Database (IEDB) ^4^ such as Bepipred ^5^, or Discotope ^6^ or others (Table 1) (^7–13^), are tools able to rapidly, in the minute-scale, generate lists of continuous and discontinuous potential B-cell epitopes. However, even state-of-the-art B-cell epitope prediction tools often output lists of potential epitopes that are too large to validate experimentally^14^. Moreover, many of the predicted epitopes will not necessarily function *in vivo* ^14^. Hence, the development of new predictive tools that enable a refinement of the computational B-cell epitope predictions available is a priority. Such tools will enable a rapid and accurate reaction in front of emergency situations such as COVID-19 pandemics or the appearance of new variants of concern that escape the immune response and derived immunotherapies ^15,16^.

To this end, we have designed Brewpitopes, a new predictive pipeline that integrates additional important features of known epitopes, such as glycosylation or structural accessibility, to curate B-cell epitope predictions for neutralizing antibody recognition and enrich that final list of potential epitopes with those that are more likely to be functional *in vivo*. We also have validated our pipeline by predicting epitopes in antibody binding regions in the proteome of SARS-CoV-2, with a special focus on the S protein and its variants of concern.

## Material and Methods

All protein three-dimensional figures have been generated with PyMol 2.5 and Chimera X. All statistical analyses have been done using R statistical software (R version 3.6.3). All data and software can be obtained from public sources for academic use.

### Dataset curation

The SARS-CoV-2 proteome in UniprotKB consists of 16 reviewed proteins ^17^. We used the corresponding FASTA sequences as starting data for linear epitope predictions. To perform structural epitope predictions, when available, we obtained the PDB structures from the Protein Data Bank database selecting the structures with the best resolution and more protein sequence coverage^18^. For those proteins with no available structure in PDB, we used Alphafold2.0 ^19^ or Modeller ^20^ to model their 3D structure.

### Linear epitope predictions

To predict linear epitopes on protein sequences we used ABCpred ^21^ and Bepipred 2.0 ^5^.

We used ABCpred ^21^, an artificial neural network trained on B-cell epitopes from the Bcipep database ^22^ to predict linear epitopes given a FASTA sequence. The identification threshold was set to 0.5 as indicated by default (accuracy 65.9% and all the window lengths were used for prediction (10-20mers). Additionally, we kept the overlapping filter on. To further augment the specificity of the predictions, we increased the ABCpred score to 0.8.

In addition, we used Bepipred 2.0 ^5^, a random forest algorithm trained on epitopes annotated from antibody-antigen complexes, as a second source to predict linear epitopes. The epitope identification threshold was set to >= 0.55 leading to a specificity of 0.81 and a sensitivity of 0.29(32).

### Structural epitope predictions

We used PDBrenum ^23^ to map the PDB residue numbers to their original positions at the UniprotKB FASTA sequence. The reason behind this step was that factors such as the inclusion of mutations to stabilize the crystal may lead to discordances between the residue numbers in the PDB and FASTA sequence from the same protein.

In order to model those SARS-CoV-2 proteins with missing structures in PDB, we used Alphafold 2.0 ^19^. We then refined the models by restraining our analysis to those regions with a pDLLT threshold of 0.7 to only assess highly confident regions. The proteins that required Alphafold modelling were M, NS6, ORF9C, ORF3D, ORF3C, NS7B and ORF3B.

To predict conformational or structural B-cell epitopes we used Discotope 2.0, a method based on surface accessibility and a novel epitope propensity score ^6^. The epitope identification threshold was set to -3.7, as specified by default, which determined a sensitivity of 0.47 and a specificity of 0.75.

### Epitope extraction and integration

Bepipred 2.0 ^5^, ABCpred ^21^ and Discotope 2.0 ^6^ predictions resulted in different tabular outputs. To extract and curate the predicted epitopes, we created a suite of computational tools in R statistical programming language and Python, available at https://github.com/rocfd/brewpitopes.

### Subcellular location predictions

When publicly available, the protein topology information was retrieved from the subcellular location section in UniprotKB ^17^. For those proteins with unavailable topology, we predicted their extraviral regions using Constrained Consensus TOPology prediction(CCTOP) ^24^, a consensus method based on the integration of HMMTOP ^25^, Membrain ^26^, Memsat-SVM ^27^, Octopus ^28^, Philius ^29^, Phobius ^30^, Pro & Prodiv ^31^, Scampi ^32^ and TMHMM ^33^. The .xml output of CCTOP was parsed using an in-house R script (xml_parser.R) and then, the extracted topology served as reference to select epitopes located in extraviral regions using the script Epitopology.R.

### Glycosylation predictions

To investigate *in-silico* which residues would be glycosylated, we used NetNGlyc 1.0 ^34^ for N-glycosylation and Net-O-Glyc 4.0 ^35^ for O-glycosylations. NetNglyc uses an artificial neural network to examine the sequences of human proteins in the context of Asn-Xaa-Ser/Thr sequons. NetOglyc produces neural network predictions of mucin type GalNAc O-glycosylation sites in mammalian proteins. We parsed the corresponding outputs using tailored R scripts and then, we extracted the glycosylated positions to filter out those epitopes containing glycosylated residues using Epiglycan.py.

### Accessibility predictions

To predict the accessibility of epitopes within their parental protein structure we computed the relative solvent accessibility (RSA) values using ICM browser from Molsoft ^36^. We used an in-house IEC browser script (Compute_ASA.icm) to compute RSA and we considered buried those residues with RSA threshold less than 0.20. Then the ICM-browser output parsed to extract the buried positions, which then served as filter to discard epitopes containing inaccessible or buried residues using Episurf.py.

### Variants of concern (VOCs) analysis

The mutations accumulated by the variants of concern Alpha, Beta, Delta, Gamma and Omicron in the S protein were obtained from CoVariants webpage ^37^, which is empowered by GISAID data ^38^. A fasta sequence embedding each variant’s mutations was generated using fasta_mutator.R.

### Data availability

All the scripts that conform the Brewpitopes pipeline can be encountered at *https://github.com/rocfd/brewpitopes* with extensive documentation.

## Results

### Brewpitopes, a pipeline to curate B-cell epitope predictions based on determinant features for *in vivo* antibody recognition

While there are some tools available to predict the presence of B-cell epitopes in a protein sequence or structure, these tools are mainly based on machine learning methods trained with experimentally validated epitopes (Table 1). However, these methods sometimes do not account for other factors that might affect the antigenicity or the potential of a protein region to be recognized specifically by antibodies.

Brewpitopes was designed as a streamlined pipeline that generates a consensus between linear and conformational epitope predictions and curates them following the *in vivo* antibody recognition constraints (Fig.1A, B, C). To this end, a suite of computational tools was created to integrate the output of different predictors of all the aforementioned features and create a filtered version of B-cell epitope regions (Fig. 2).

**Figure 1A.**
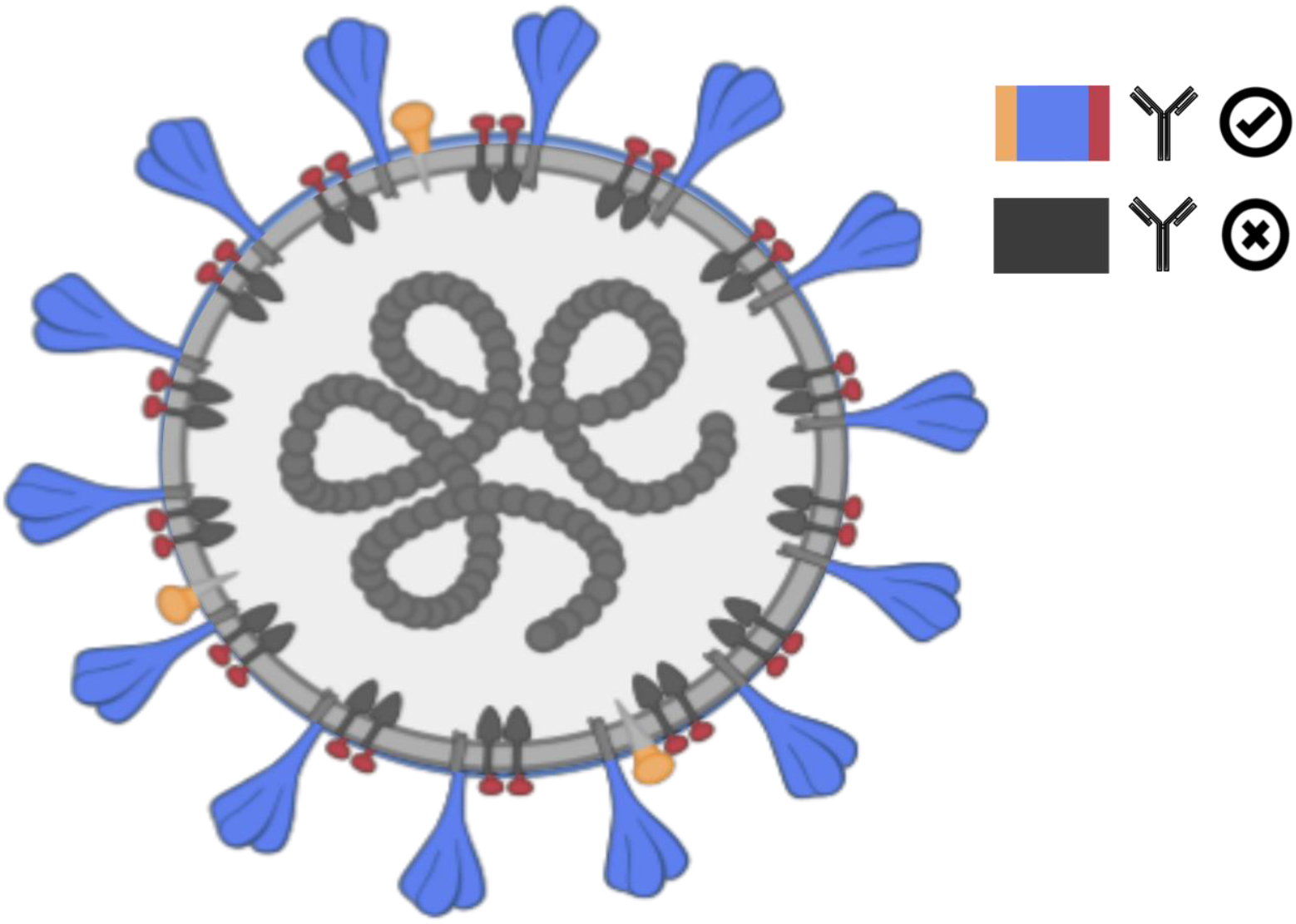
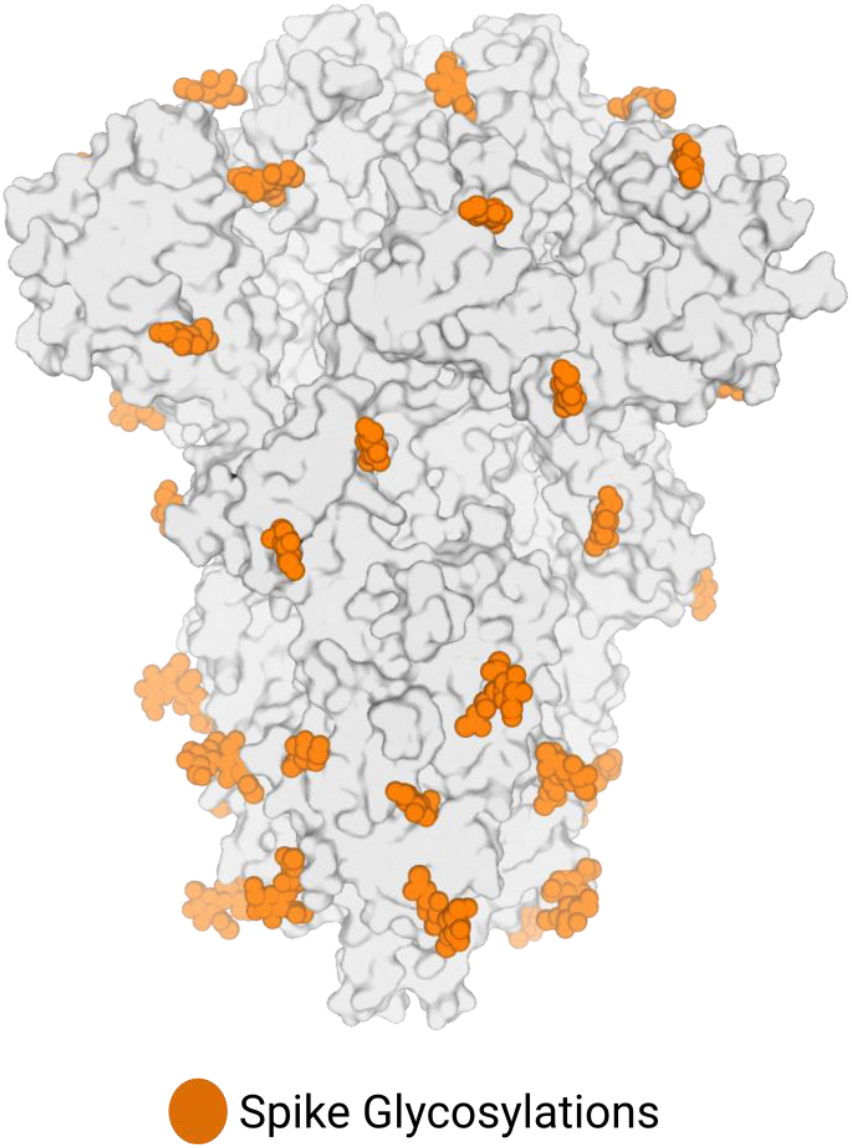
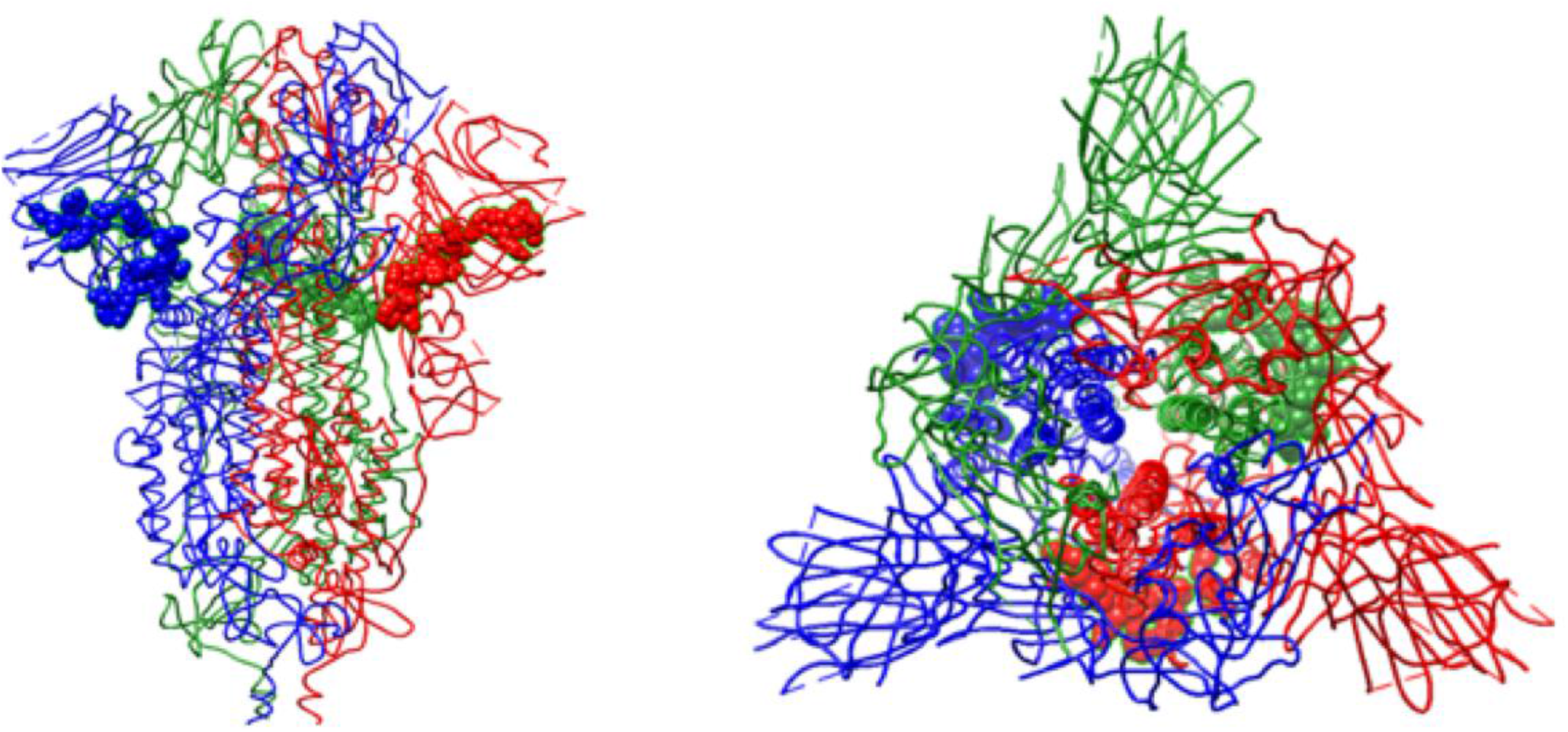
Biophysical constraints for *in vivo* antibody recognition. A) Recognition of extraviral protein regions. Neutralizing antibodies can only inspect the external surface of viral particles. Hence, only epitopes located in extraviral regions will be recognizable by this type of antibodies. In Brewpitopes, we used protein topology information or predictions to assess the subcellular location of the analyzed protein regions. Once defined, we only selected those epitopes located on extracellular protein regions. Hence, epitopes located in transmembrane or intracellular protein regions were discarded. B) Glycan coverage prevents *in vivo* antibody recognition of an epitope. Epitopes that contain glycosylated residues will be covered by the glycan and therefore shoud be discarded as candidates for antibody recognition. In Brewpitopes, to predict the glycosylation profiles of target proteins we used Net-N-glyc and Net-O-glyc to predict N- and O-glycosylations respectively. Once predicted, we selected those epitopes that do not contain any glycosylated position. C) Epitope accessibility within the parental protein 3D structure. Epitopes that contain buried residues will not be accessible for *in vivo* antibody recognition. Left; structure of the S protein highlighting a fully exposed epitope. Right; structure of the S protein displaying a buried epitope. In Brewpitopes, we calculated the residue solvent accessibility (RSA) using the Molsoft ICM Browser software. Once predicted, we selected those epitopes with all residues over a RSA value of 0.2. These were considered the accessible epitopes whereas epitopes with a single residue with a lower RSA value than 0.2 were considered buried and therefore were discarded.

**Figure 2.**
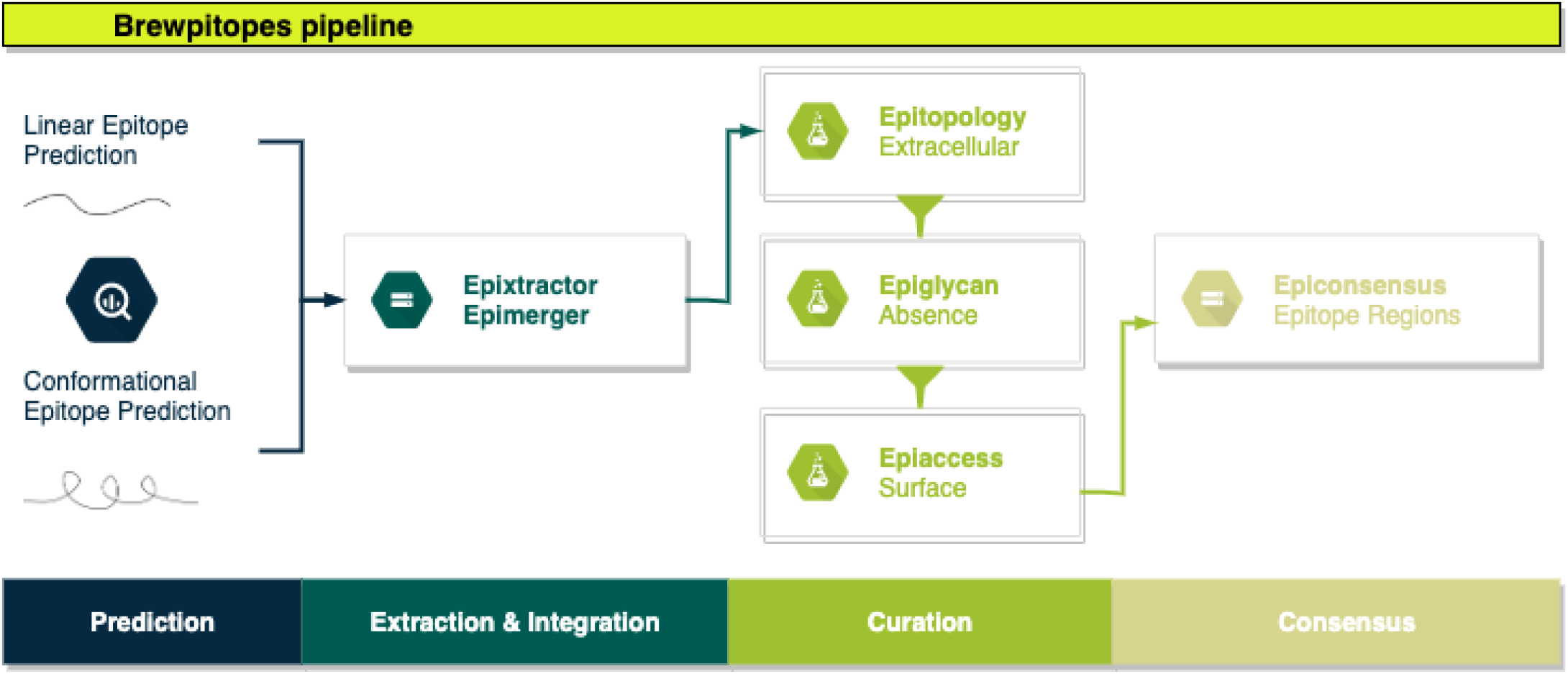
Brewpitopes pipeline to refine B-cell epitope predictions into epitopes optimized for *in vivo* antibody recognition. Linear and structural epitope predictions were obtained using Bepipred 2.0, ABCpred and Discotope 2.0 respectivelly. Then the epitopes were extracted and integrated using Epixtractor and Epimerger. Followingly, the epitopes were refined using the biophysical constraints for *in vivo* antibody recogntition (described in Figure 1). The protein topology analysis was performed using CCTOP and the epitopes located in extratraviral regions were labeled using Epitopology. The glycosylation profile was predicted using Net-N-Glyc and Net-O-Glyc and the glycosylated epitopes were labeled using Epiglycan. The accessibility analysis was computed with Molsoft taking an RSA threshold of 0.2 for buried residues. Epitopes containing buried residues were labeled using Episurf. To filter the candidate epitopes according to the biophysical constraints (extraviral location, absence of glycosylation coverage and accessibility within the 3D protein structure) we used Epifilter. Finally, since many predicted epitopes were overlapping due to their prediction using different tools, they were converged into epitope regions using Epiconsensus.

Linear epitopes are continual stretches of residues located at the surface of proteins whereas conformational epitopes are discontinuous residues recognized due to their structural disposition. For both cases exist state-of-the-art predictors (Table 1). To start with, in Brewpitopes, we have predicted linear epitopes using Bepipred 2.0 ^5^ and ABCpred^21^ and we have searched for conformational epitopes using Discotope 2.0 ^6^. Once predicted, we have extracted the epitopes using tailored R scripts named Epixtractor and we then integrated the results using Epimerger.

First, since neutralizing antibodies can only inspect the external surface of cells or viral particles, we propose that those epitopes predicted in intracellular and transmembrane regions of proteins cannot be targeted by these type of antibodies (Fig. 1A). Hence, the subcellular location of an epitope is a recognition constraint ^39^, which our pipeline uses to prioritize epitopes located on extracellular protein regions while discarding those located in intracellular and transmembrane regions. To predict the subcellular location of a protein region; we used protein topology information. For some proteins, the topology is already available at UniprotKB ^17^, however, for some others it is not. In such cases, the alternative is to predict the topology of the protein. When this occurred, we used CCTOP to predict their transmembrane, intracellular and extracellular regions ^24^. Once we had the extracellular regions, we labelled the epitopes using Epitopology.

Next, glycans can cover B-cell epitopes, limiting their accessibility for antibody recognition (Fig. 1B) ^40^. For this reason, our pipeline uses *in-silico* tools to predict glycosylated sites and, discards all the epitopes overlapping glycosylated residues. Concretely, we have used NetNglyc1.0 ^34^ and NetOglyc4.0 ^35^, for the prediction of N-glycosylations and O-glycosylations respectively. These methods are based on artificial neural networks trained on glycosylation patterns by which they can predict glycosylation sites *ab initio* given a protein sequence. Once the glycosylated positions were retrieved, we used Epiglycan to label epitopes with one or more glycosylations as “Glycosylated” and those with no glycans as “Non-glycosylated”.

The accessibility of the epitope within the protein structure is another antibody recognition constraint ^41^ (Fig 1C). Accordingly, our pipeline calculates the relative solvent accessibility (RSA) values for all the residues and filters out those epitopes containing buried amino acids. To compute the RSA values based on crystal structures we have used Molsoft ^36^ and the *in-house* script compute_asa.icm. Overall, we have determined the buried positions as those residues with RSA values lower than 0.20. Next, these positions served as reference to label epitopes containing at least one buried residue as “buried”, which therefore would not be accessible for antibody recognition. Conversely, epitopes with all the residues calculated as exposed were labelled as “accessible”.

Finally, to conclude the Brewpitopes pipeline we used Epifilter to filter out those epitopes that were labelled as intracellular, glycosylated or buried. Additionally, a length filter was used to select epitopes longer than 5 amino acids, since shorter peptides were considered unspecific. Therefore, our final candidates were extracellular, non-glycosylated and accessible enhancing their capacity to be recognized by antibodies *in vivo*. In order to merge overlapping epitopes into epitope regions and to generate a consensus between B-cell epitope predictors, we designed Epiconsensus; a tool that not only merges overlapping epitopes but it also scores the resulting epitope regions with the following an ordering criteria between predictors: first, ordered by best Bepipred 2.0 score ^5^; second, ordered by best Discotope 2.0 score ^6^ and third, ordered by best ABCpred score ^21^.

### Bioinformatic validation of Brewpitopes in the proteome of SARS-CoV-2

Brewpitopes can be implemented to any target protein or organism, but due to the pandemic context and the interest in B-cell epitopes and neutralizing antibodies against SARS-CoV-2, to validate the pipeline we analysed its proteome. Within SARS-CoV-2, we specially focused on the S protein due to its importance in vaccine design, therapeutic antibodies and immune evasion ^42^. Our results confirm the neutralizing potential of the S protein and also show how other proteins of SARS-CoV-2 contain other regions with potential epitopes for neutralizing antibodies.

Focusing on the S protein, linear epitope predictions resulted in 213 epitopes and structural predictions in 6. Once integrated, 10 epitopes were discarded due to their intraviral location. Next, since it had been established that S protein is heavily glycosylated (15), 52 epitopes were filtered out due to the presence of glycans. Lastly, 143 epitopes were discarded because they contained at least one residue buried within the 3D structure of the S protein. As a result, 14 epitopes derived from S were curated for optimized antibody recognition (Fig.3). Compared to the initial state-of-the-art epitope predictions, our results show that only a 5.5% of the predicted epitopes for the S protein will be enriched for antibody recognition *in vivo* due to the recognition constraints we have analysed with Brewpitopes (Fig.4). Furthermore, to generate a consensus between linear and conformational predictions from different tools, the overlapping epitopes were merged into epitope regions. In the case of the S protein, the 14 candidates were merged into 7 epitope regions (Fig 3 & Table 4).

**Figure 3.**
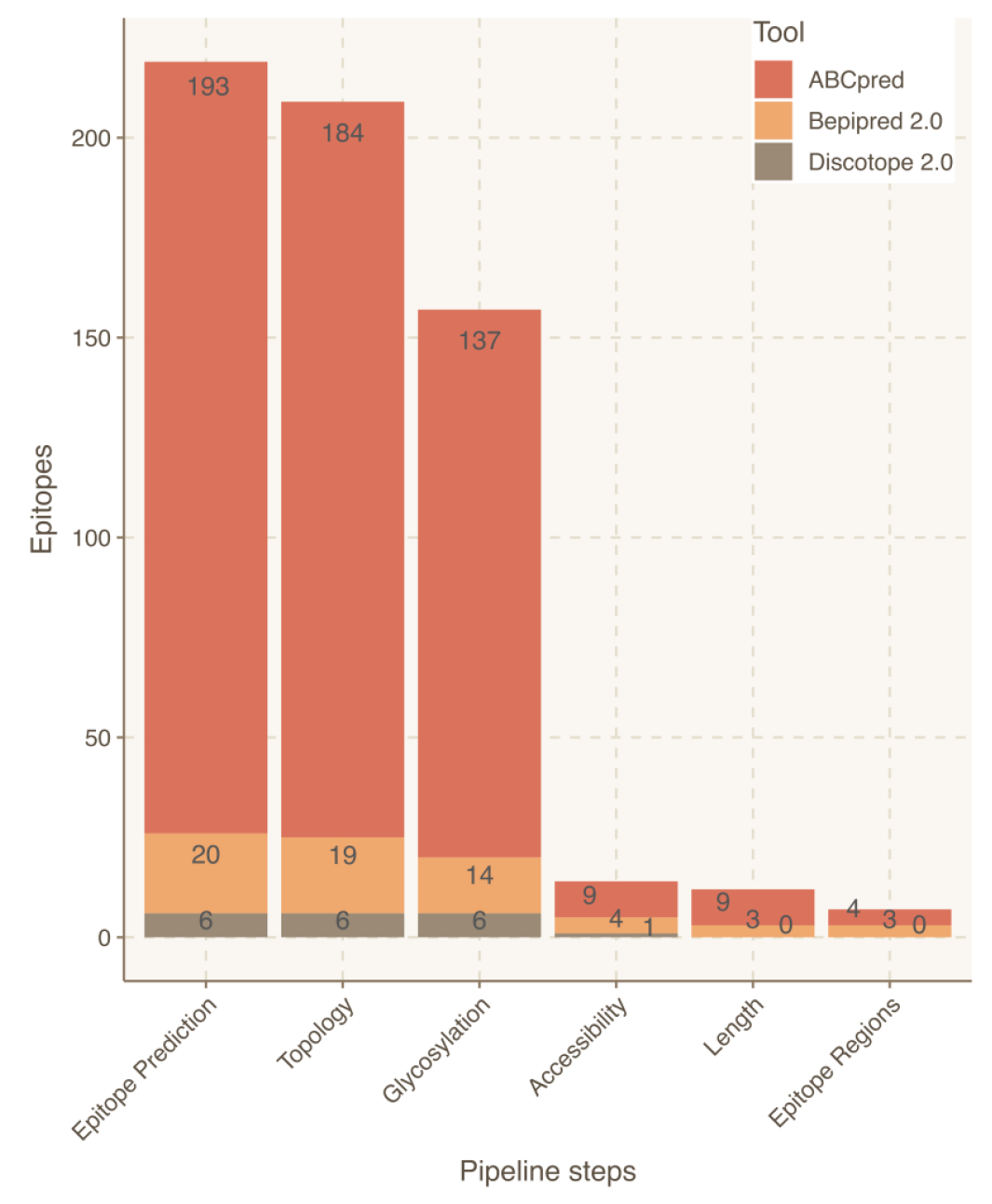
Epitope refinement for the WT S protein using Brewpitopes. The X-axis represents the steps of the Brewpitopes pipeline and the Y-axis the number of epitopes selected by each filtering step of Brewpitopes (Fig.2). The length filter was used to select epitopes longer than r or more amino acids, since shorter peptides were considered inespecific.

**Figure 4.**
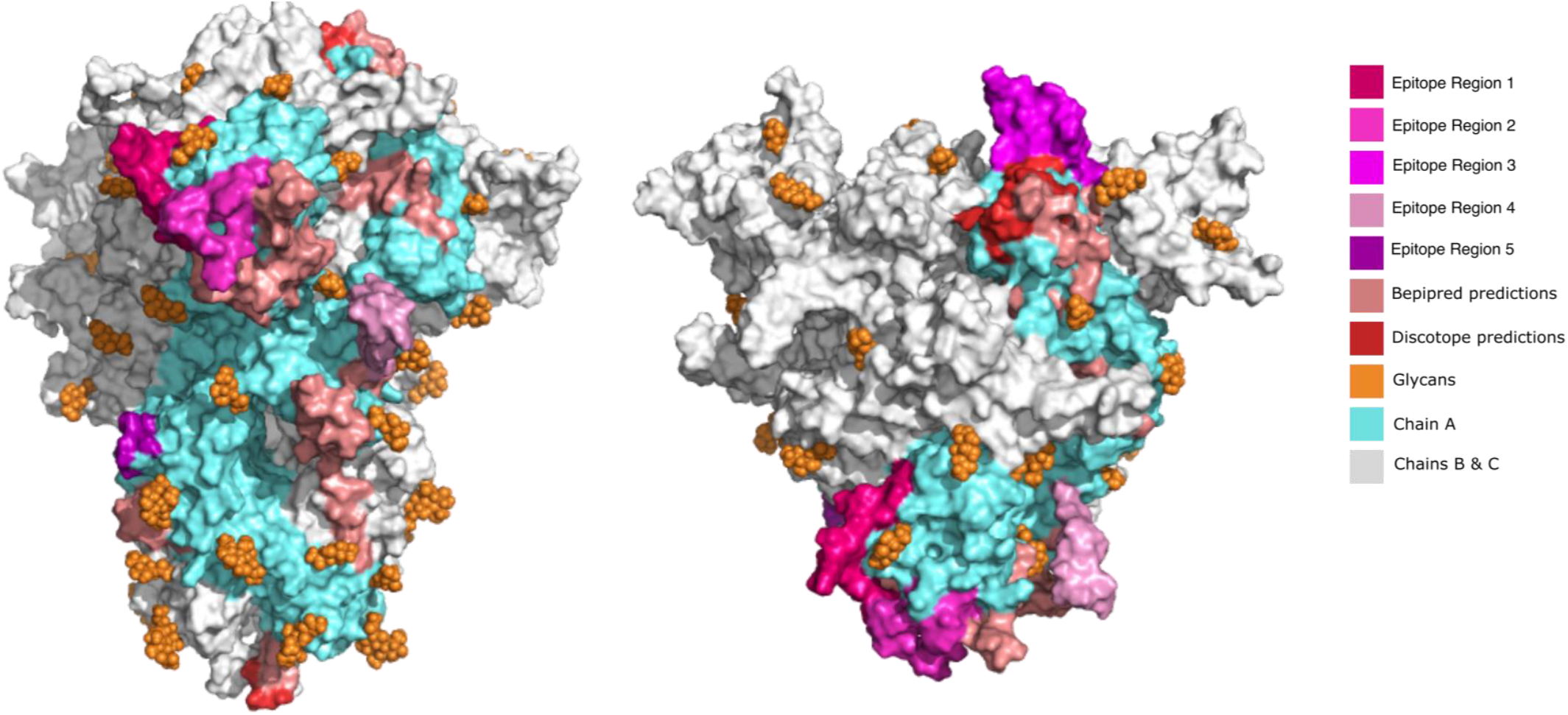
Epitope location on the 3D structure of the S protein. Left) Front view of the S protein 3D structure. Right) Top view. All the epitopes were only labelled on the chain A of the S protein for visualization purposes (blue). The epitope regions 6 and 7 were not displayed because they escaped the limits of the PDB structure. Also, the great number of predictions by the software ABCpred precluded their 3D representation. In this figure, it can be observed how the region of initial predictions is refined into smaller regions represented by the curated epitopes.

As an external validation, the epitope regions identified in the S protein were cross-validated with the epitopes described at the IEDB database ^43^ (Table 4). Notably, the regions identified in our pipeline were all encountered among IEDB epitopes, which confirms the validity of our predictions. However, our epitope regions represented less than 1% of the epitopes for the S protein listed in the IEDB. Compared to the initial output from the computational tools, the final list of prioritized epitopes from our pipeline was enriched 5-fold in validated epitopes from IEDB (p < 2e-4). This confirms the power of Brewpitopes to refine B-cell epitope computational predictions to a reduced set of epitopes with greater probability for *in vivo* antibody recognition.

To extend our analysis to the rest of the SARS-CoV-2 proteins in search of other epitopes with antibody recognition potential, we used Brewpitopes to analyse full proteome of the virus. Overall, 4/15 of the remaining proteins contained candidate epitopes for neutralizing antibodies (R1AB, R1A, AP3A & ORF9C) (Table 3). The reason behind the remaining proteins (11/15) not containing epitopes on antibody recognition sites was mainly their major intraviral location (NS7A, NS7B, ORF3D, ORF3C, ORF9B, ORF3B, NS8, NS6, M, E & N) and the absence of epitopes in their extraviral regions. Within the proteins that contained candidate epitopes, R1AB and R1A stood out carrying 479 and 348 epitopes respectively. Such high numbers of epitopes can be mainly explained due to their long sequences, 7096 and 4405 amino acids respectively. Remarkably, R1A corresponds to the N-terminal region of R1AB so many of the predicted epitopes are shared. Furthermore, R1AB is a polyprotein cleaved int o 15 chains that here were analysed altogether using the Uniprot entry. Differently, ORF9C (4) and AP3A (2) presented a lower number of epitopes. In terms of epitope regions, R1AB counted with 62 regions, R1A with 46, ORF9C with 2 and AP3A with 1. Altogether these results corroborate that four proteins other than S have at least one epitope region candidate for *in vivo* antibody recognition.

**Table 3.**
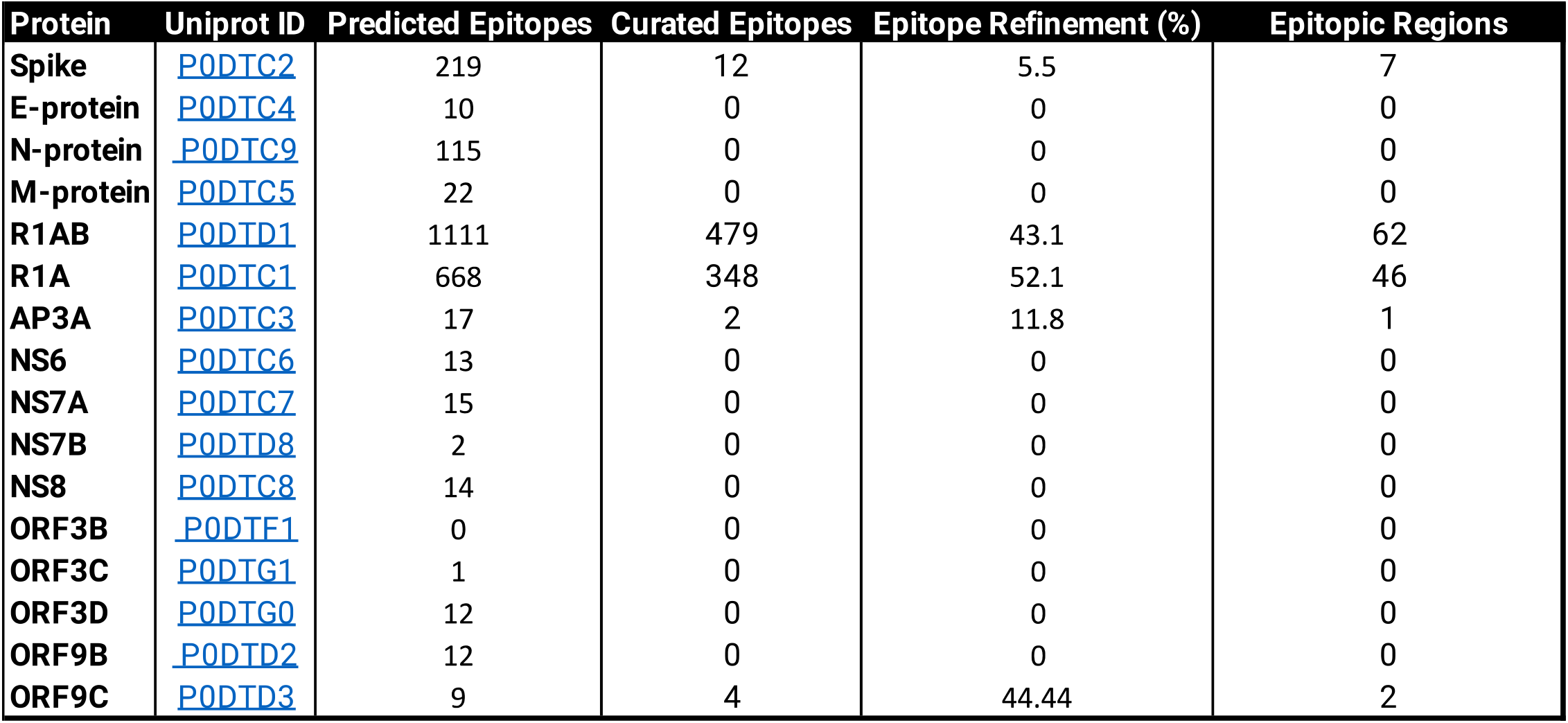
Epitope refinement for the SARS-CoV-2 proteome using Brewpitopes. Predicted epitopes correspond to the number of epitopes obtained using linear and structural predictions. Curated epitopes refers to the refined epitopes obtained using Brewpitopes. Epitope refinement is the percentage of curated epitopes over predicted epitopes. The epitope regions are the result of converging overlapping epitopes using Epiconsensus.

| Protein | Uniprot ID | Predicted Epitopes | Curated Epitopes | Epitope Refinement (%) | Epitopic Regions |
| --- | --- | --- | --- | --- | --- |
| **Spike** | P0DTC2 | 219 | 12 | 5.5 | 7 |
| **E-protein** | P0DTC4 | 10 | 0 | 0 | 0 |
| **N-protein** | P0DTC9 | 115 | 0 | 0 | 0 |
| **M-protein** | P0DTC5 | 22 | 0 | 0 | 0 |
| **R1AB** | P0DTD1 | 1111 | 479 | 43.1 | 62 |
| **R1A** | P0DTC1 | 668 | 348 | 52.1 | 46 |
| **AP3A** | P0DTC3 | 17 | 2 | 11.8 | 1 |
| **NS6** | P0DTC6 | 13 | 0 | 0 | 0 |
| **NS7A** | P0DTC7 | 15 | 0 | 0 | 0 |
| **NS7B** | P0DTD8 | 2 | 0 | 0 | 0 |
| **NS8** | P0DTC8 | 14 | 0 | 0 | 0 |
| **ORF3B** | P0DTF1 | 0 | 0 | 0 | 0 |
| **ORF3C** | P0DTG1 | 1 | 0 | 0 | 0 |
| **ORF3D** | P0DTG0 | 12 | 0 | 0 | 0 |
| **ORF9B** | P0DTD2 | 12 | 0 | 0 | 0 |
| **ORF9C** | P0DTD3 | 9 | 4 | 44.44 | 2 |

**Table 4.** Epitope crossvalidation with the IEDB database. The enrichment of the epitope regions identified by Brewpitopes were compared versus the total epitopes validated in the IEDB database for the S protein of SARS-CoV-2. This lead to a Fisher’s exact test p-value <= 0.002065 and a five-fold enrichment in the odds ratio.

|  | Fisher's P-value | Odds Ratio |
| --- | --- | --- |
| **Brewpitopes** | 0.0002065 | 4.995633 |

### Analysis of epitope conservation in the S protein of SARS-Cov-2’s Variants of Concern

To study whether the mutations accumulated in the S protein of the variants of concern (Alpha, Beta, Delta, Gamma and Omicron) could lead to immune escape, we used the Brewpitopes pipeline to analyse the different variants. We generated tailored FASTA files for the mutations of each variant and we retrieved the structures from PDB when available. For the Omicron variant, we modelled its structure using Modeller ^20^. Once we had run Brewpitopes, we compared the final number of epitopes with neutralizing potential per each variant with the epitopes generated by our analysis of the Wuhan S protein, considered the wildtype. Concretely, we aimed at identifying epitope losses due to the presence of mutations, new glycosylations and newly buried positions. Additionally, we also looked for new epitopes generated by the unique mutations of each variant. To compare the epitope regions in the WT variant versus those of the variants of concern (VOCs), the length of these epitope regions was added and divided by the length of the S protein to obtain the epitope coverage (i.e., the fraction of the protein sequence that is covered by epitopes). These lead to an epitope coverage of 9.43% for the WT variant, a value that will be used as reference to assess the epitope conservation on the VOCs.

To visualize the accumulation of mutations in the S protein in the VOCs, we calculated the intersections of shared mutations between variants (Table 5 and Fig.5). As it can be observed in the upset plot, Omicron variant accumulates the most mutations (37), of which 28 are exclusive for this variant. Gamma accumulates 8 unique mutations, Delta 7, Beta 6 and Alpha 4. Also, the degree of shared mutations is low being Alpha and Omicron the variants that share more mutations, with 4. The other combinations only share a single mutation. Finally, the intersection of all variants only leads to a single common mutation. This high diversity in the mutations accumulated in S points towards separate evolution paths and can derive to variant-specific mechanisms of immune evasion and decreased antibody recognition. The fact that Omicron accumulates more than 3-times more mutations at S than the remaining VOCs indicates a greater potential for epitope disruption.

**Figure 5.**
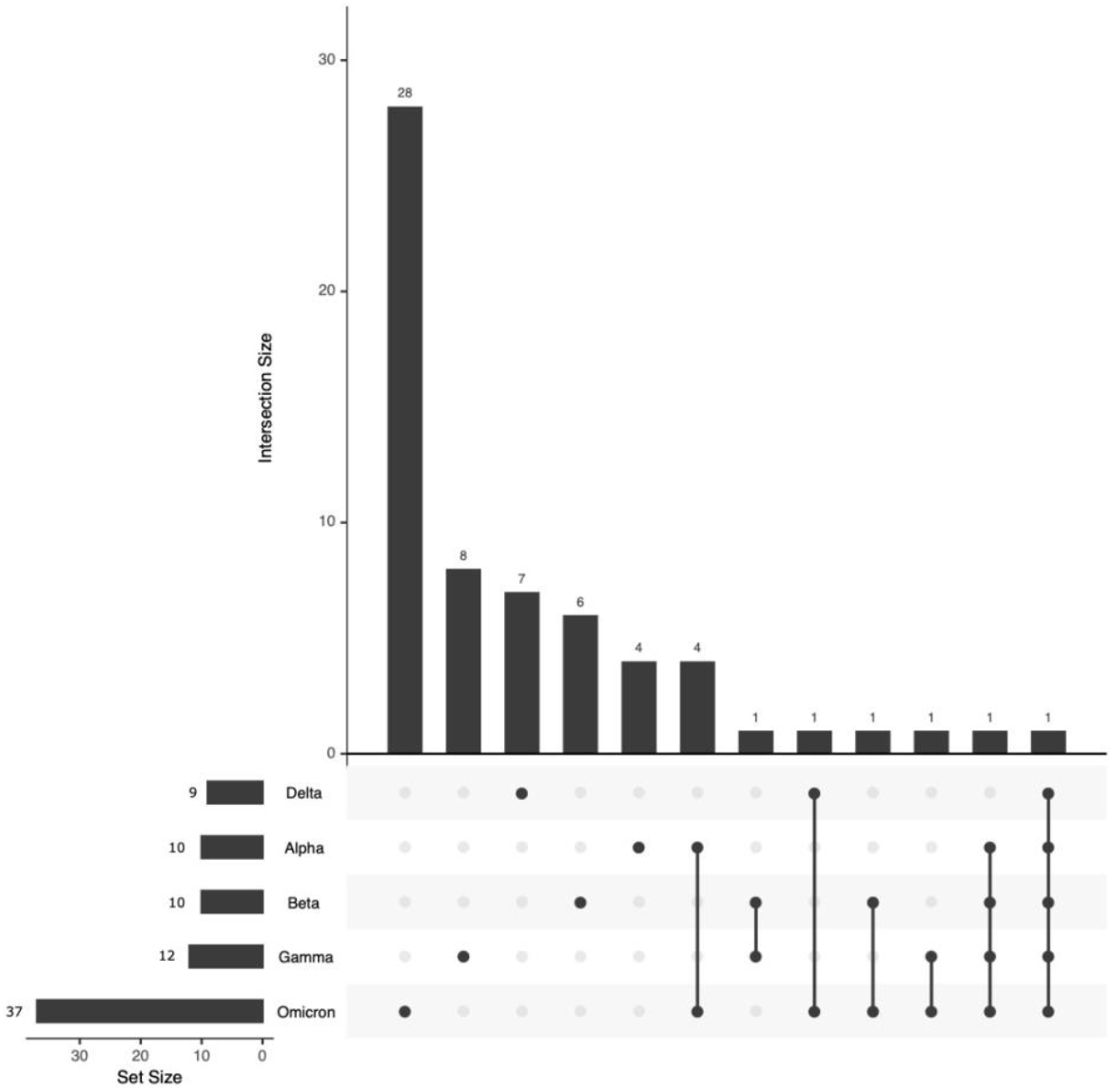
Mutations accumulated in the protein S of the Variants of Concern Alpha, Beta, Delta, Gamma and Omicron. The upset plot represents the unique mutations accumulated in each VOC and the mutations shared by two or more variants. The total mutations per each variant are displayed in the lower barplot. The accumulation of mutations in the S protein can be linked to a greater potential of immune escape due to the protential disruption of epitopes by the mutations. The variant Omicron stands out with the 3 times more mutations than the other variants and has the greatest potential for immune escape.

**Figure 6.**
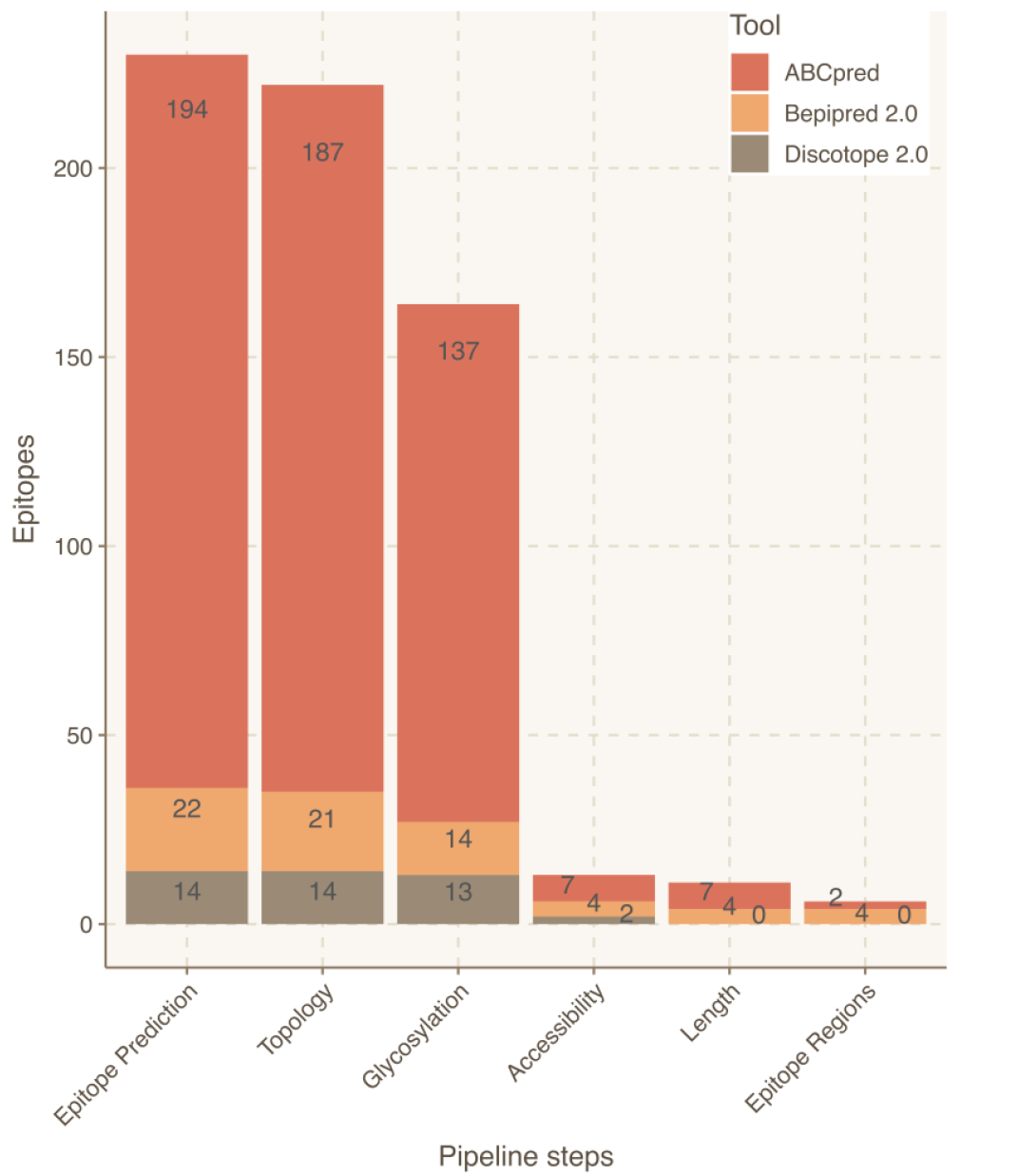
Epitope refinement for the S protein of the Omicron variant using Brewpitopes.. The X-axis represents the steps of the Brewpitopes pipeline and the Y-axis the number of epitopes selected by each filtering step of Brewpitopes (Fig.2). The length filter was used to select epitopes longer than r or more amino acids, since shorter peptides were considered inespecific.The epitpoe yield obtained with the Omicron variant can be compared to the WT’s yield in Figure 3.

Comparatively, each VOCs accumulated specific mutations in the S protein (Table 5). These lead to a specific epitope landscape per variant (Tables 3 and 6) that differed to the observed in the Wuhan variable, considered the WT. Considering epitope regions, Alpha loses the ER7; Beta loses ER4 and ER7 but gains an epitope region at 828-845; Gamma loses ER2, ER3 and ER4; Delta loses ER3, ER4 and ER6 but gains ER1, ER5 and ER8 and Omicron loses partially ER2, ER3 and ER4 and entirely ER7 (Table 7).

In terms of epitope coverage, the major loss occurs in Gamma (4%) and Omicron (2%) while Alpha and Beta lose less than 1.5%. Differently, Delta gains a 0.5% in epitope coverage in respect to the Wuhan due to large epitope gains (Table 4). The differences in epitope landscape in the VOCs can indicate partial losses in antibody recognition. However, using Brewpitopes, a core of epitope regions recognized across variants was encountered (Table 7).

**Table 5.** Mutations of the VOCs in the S protein.

| **VOC** | **Mutations** |
| --- | --- |
| **Alpha** | H69-, V70-, Y144-, N501Y, A570D, D614G, P681H, T716I, S982A and D1118H |
| **Beta** | D80A, D215G, L241-, L242-, A243-, K417N, E484K, N501Y, D614G and A701V |
| **Gamma** | L18F, T20N, P26S, D138Y, R190S, K417T, E484K, N501Y, D614G, H655Y, T1027I and V1176F |
| **Delta** | T19R, E156-, F157-, R158G, L452R, T478K, D614G, P681R and D950N |
| **Omicron** | AG7V, H69-, V70-, T95I, G142-, V143-, Y144- Y145D, N211-, L212I, G339D, S371L, S373P, S375F, K417N, N440K, G446S, S477N, T478K, E484A, Q493R, Q496S, Q498R, N501Y, Y505H, T547K, D614G, H655Y, N679K, P681H, N764K, D796Y, N856K, Q954H, N969K, L981F, 214EPE |

**Table 6.** Results of the epitope refinement of Brewpitopes on the protein S of the Variants of Concern Alpha, Beta, Delta, Gamma and Omicron. Predicted epitopes correspond to the number of epitopes obtained using linear and structural predictions. Curated epitopes refers to the refined epitopes obtained using Brewpitopes. The epitope regions are the result of converging overlapping epitopes. Epitope refinement is the percentage of curated epitopes over predicted epitopes. Epitope conservation refers to the percentage of epitopes refined in the variant that are also indentified in the WT S protein. Epitope region conservation refers to the percentage of epitope regions shared between the variant and the WT S protein.

| Variant | ID | Predicted Epitopes | Curated Epitopes | Epitopic Regions | Epitope Refinement (%) | Epitope Conservation (%) | Epitopic Region Conservation (%) |
| --- | --- | --- | --- | --- | --- | --- | --- |
| **Wuhan-2** | WT | 219 | 12 | 7 | 5.5 | 100.0 | 100.0 |
| **Alpha** | B.1.1.7 | 206 | 13 | 6 | 6.3 | 108.3 | 85.7 |
| **Beta** | B.1.351 | 225 | 11 | 6 | 4.9 | 91.7 | 85.7 |
| **Delta** | P.1 | 213 | 15 | 7 | 7.0 | 125.0 | 100.0 |
| **Gamma** | B.1.617.2 | 214 | 6 | 5 | 2.8 | 50.0 | 71.4 |
| **Omicron** | B.1.1.529 | 230 | 11 | 6 | 4.8 | 91.7 | 85.7 |

**Table 7.**
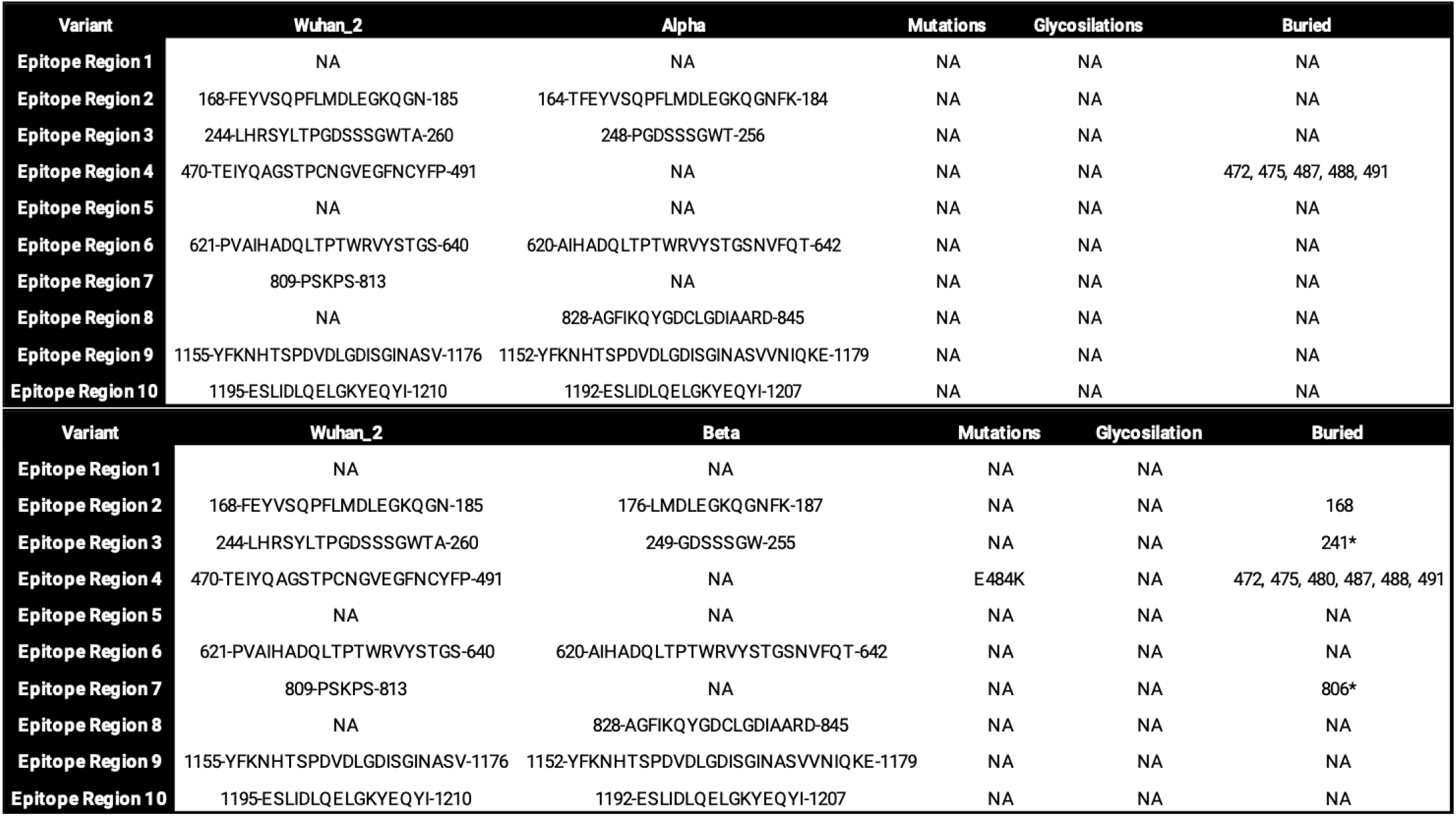
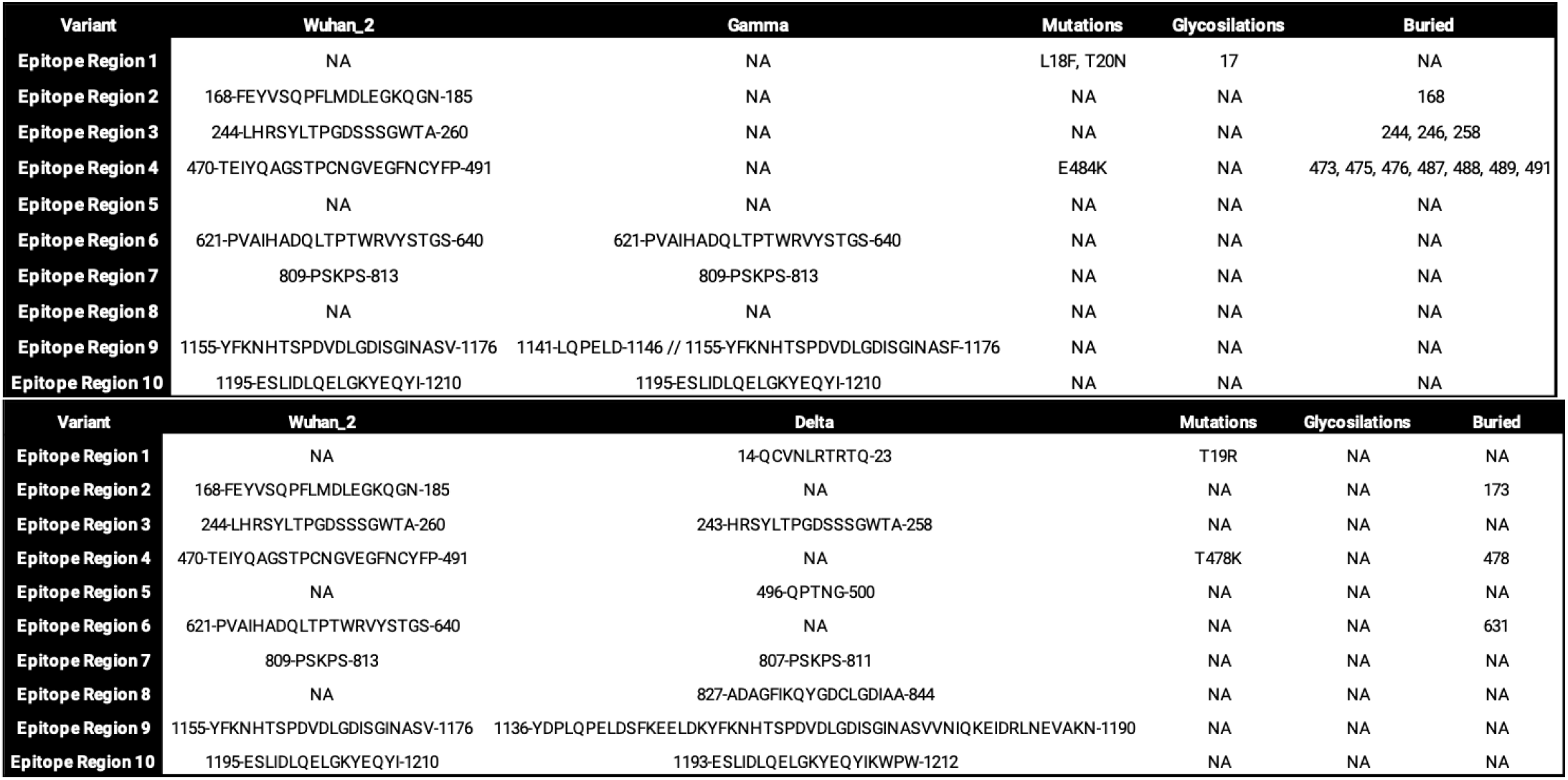
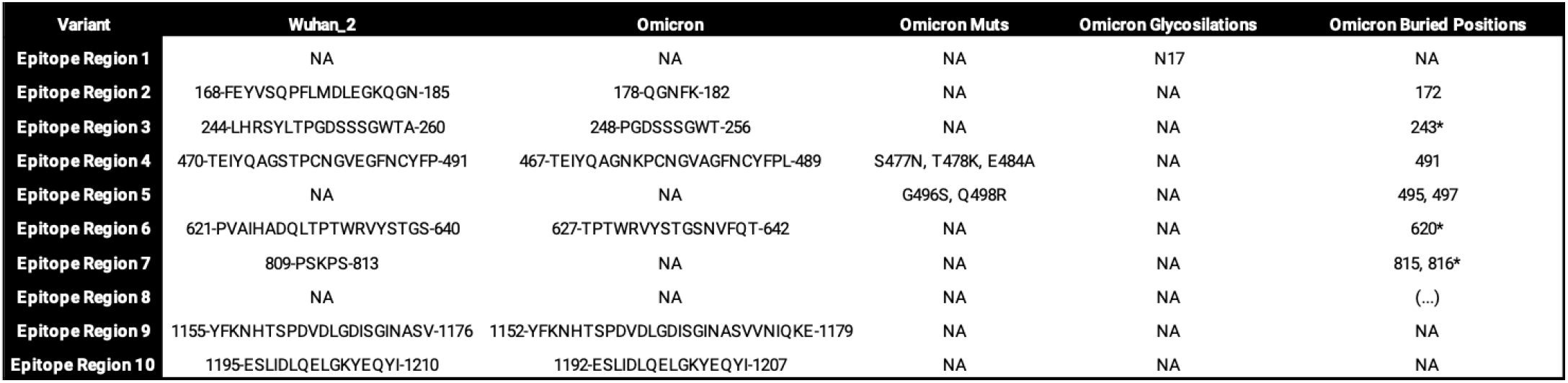
Epitope regions identified in the WT S protein using Brewpitopes compared to the epitope regions of the Variants of Concern. Mutations, Glycosylation and Buried refer to the residues that are causing the disappearance of epitope regions in the Variants of Concern.

## Discussion

*In vivo* antibody recognition is constrained by molecular features not often integrated altogether in state-of-the-art B-cell epitope predictors, including the extracellular location of the epitope, the absence of glycosylation coverage and the surface accessibility within the parental protein structure (Table 1). We have integrated them into Brewpitopes, a pipeline for the refinement of B-cell epitope predictions optimized for *in vivo* antibody recognition. The pipeline has been used to analyse the proteome of SARS-CoV-2 resulting in a refined set of epitopes with neutralizing potential located, not only on the S protein and its VOCs, but also in four additional proteins. As exemplified in the SARS-CoV-2 analysis, Brewpitopes is a ready-to-use tool that will enhance the accuracy and throughput of B-cell epitope predictions for future public health emergencies such as new COVID-19 variants or other potential pathogenic threads.

The B-cell epitope profiling in SARS-CoV-2 proteins has been a research-intensive topic since the start of the COVID-19 pandemic due to its implications for the development of vaccines and therapeutic antibodies (Table 2) ^2,44–53^. However, as shown in tables 1 and 2 none of the proposed strategies integrates the three aforementioned features altogether. In the same line, there is no available bioinformatics pipeline with a streamlined implementation of the biophysical constraints of the epitope recognition microenvironment as we have presented here with Brewpitopes. Furthermore, the available methods predict a single type of epitope, be it linear or conformational, whereas with Brewpitopes we propose an integration of both with our Epiconsensus tool. All these ponder Brewpitopes as a unique tool for the refinement of B-cell epitope predictions into epitopes with increased neutralizing potential. Overall, the epitope landscape of SARS-CoV-2 is being unravelled identifying the actionable epitope regions available for adequately adjusting the current and future preventive therapies.

Regarding the filters implemented in Brewpitopes, it should be considered that they are based on predictions, such as CCTOP for the subcellular location of protein regions or Net-N-glyc and Net-O-glyc for glycosylations. This comes with the advantage of enabling *ab initio* predictions, which do not require new experimental validations that would significantly extend the validation period and at the same time rely on the precision of the individual methods. However, relying on predictions in some cases may lead to false positives. In the case of glycosylations, we only consider the predicted glycosylated position when glycans can be large structures that could also affect the accessibility of neighbouring positions. Still which glycan occupies a position can not be predicted with in-silico methods based on the sequence and therefore this limits the estimation of which neighbouring residues could be affected. In terms of structural accessibility, we observed that some predicted epitopes had residues buried on their parental protein structure and hence, their accessibility would be difficult ^39^. Our filter discarded any epitope containing at least one buried residue. This appeared to be the most stringent filter since it downsized the number of epitopes from 137 to 14 in the case of S (Table 3). Still, unmodeled regions, not present in the crystal structure, were considered as exposed regions due to their high mobility. However, even though some of the epitopes discarded may still have antigenic activity, the resulting epitopes from Brewpitopes are expected to be more enriched immunogenically than the discarded ones. In terms of software flexibility, Brewpitopes has been built upon Discotope 2.0, and Bepipred 2.0, which during the pipeline development were considered state-of-the-art by the IEDB analysis resource tool^43^. However, Brewpitopes will be further developed to integrate more methods to keep up with the fast evolution pace of the field. Hence, maintaining state-of-the-art capacities.

While Brewpitopes can be applied to any protein or organism, given the wealth of data and biomedical interest, we decided to focus our use case on the analysis of SARS-CoV-2. In our proteome-wide analysis of epitopes with neutralizing potential, we have specially focused on the S protein and its VOCs. This led to the discovery of 6 epitope regions that are conserved across variants, which could explain the conserved antibody recognition of vaccinated patients against new variants ^54^. In this line, the restrictive nature of Brewpitopes leads to a significant reduction on the S protein epitopes predicted, which is an added value in terms of their experimental validation. In addition, our findings highlight the importance of epitopes located in other SARS-CoV-2 less variable proteins (R1AB, R1A, AP3A and ORF9C) would overcome the limitations pondered by the variants of concern. Despite these proteins are not considered structural for the virus, they contain regions predicted as extracellular which could therefore lead to accessible epitope regions.

The S protein is under very strong evolutionary pressure by the human immune system and hence, a selective advantage for those mutations that lead to decreased antibody recognition is expectable ^55,56^. For this reason, the study of the epitopes located in antibody binding regions and how mutations change these epitopes is a major public health interest. We compared the highly immunogenic epitopes predicted in the reference S protein versus those predicted on the variants Alpha, Beta, Delta, Gamma and Omicron (Table 8). Relevantly, we have observed how epitope coverage losses occur in the different variants as reported in Table 4: Gamma (4%), Omicron (2%), Beta and Alpha (1%). In contrast, Delta leads to an increased epitope coverage of 1% due to the presence of a long epitope region at 1136-1190. Overall, this analysis enabled the observation of a decreased epitope coverage in all the variants but Delta. These epitope losses indicate the variants are already evolving towards immune escape.

**Table 8.** Epitope coverage of the WT S protein versus the Variants of Concern. The epitope coverage is the percentage of the total protein sequence (the S protein) that is covered by epitope regions. It estimates the antigenicity potential of a protein. The loss of epitope coverage in the variants of concern is a proxy to estimate their immune escape potential due to the loss of *in vivo* antibody recogntition sites.

| Variant | Epitope Coverage (%) |
| --- | --- |
| Wuhan_2 | 9.43 |
| Alpha | 9.03 |
| Beta | 8.17 |
| Delta | 10.13 |
| Gamma | 5.42 |
| Omicron | 7.62 |

However, there is also a high degree of epitope conservation observed across the variants. This indicates that a core epitope group is maintained, which has beneficial implications for the efficacy of vaccines towards the variants of concern. In contrast, previous studies showed a decrease of neutralization against the Omicron variant ^57^. In our analysis, Omicron loses a 2% of epitope coverage which might partially explain the loss of neutralization, since our refinement is stringent. Discordances between previous studies ^57^ and our data might be explained both for the high stringency of our filters and by the fact that both studies followed different approaches. As aforementioned, our method can discard few antigenic epitopes not necessarily representing a true neutralization loss in vaccinated patients.

In summary, Brewpitopes is a B-cell epitope refining pipeline that can be implemented in any target protein or organism and integrates relevant features for the antibody recognition microenvironment such as glycosylation, subcellular location and surface accessibility. Furthermore, the implementation of Brewpitopes to the proteome of SARS-CoV-2 has identified epitopes that escape the mutations of the main variants of concern.

## Abbreviations

Brewpitopes: pipeline to refine B-cell epitope predictions
S protein: Spike protein of SARS-CoV-2
VOCs: Variants of Concern from SARS-CoV-2
RSA: relative solvent accessibility
IEDB: ImmunoEpitope Database
Alpha, Beta, Delta, Gamma & Omicron: SARS-CoV-2 Variants of Concern.

---

**Supplementary Table 1.** Glycosylation predictions Vs MS-validated glycosylation sites. Predictions were performed using Net-N-Glyc for N-glycosylations and Net-O-glyc for O-glycosylations.

| MS-validated Glycosylation sites | Reference | Predicted Glycosylation sites |
| --- | --- | --- |
| N17 | Watanabe et al., Zhang et al. | Yes |
| N16 | Watanabe et al., Zhang et al. | Yes |
| N74 | Shajahan et al., Zhang et al., Watanabe et al. | Yes |
| N122 | Watanabe et al., Zhang et al. | Yes |
| M149 | Shajahan et al., Zhang et al., Watanabe et al. | Yes |
| N165 | Shajahan et al., Zhang et al., Watanabe et al. | Yes |
| N234 | Shajahan et al., Zhang et al., Watanabe et al. | Yes |
| N282 | Shajahan et al., Zhang et al., Watanabe et al. | Yes |
| N331 | Watanabe et al., Zhang et al. | Yes |
| N343 | Watanabe et al., Zhang et al. | Yes |
| N603 | Watanabe et al., Zhang et al. | Yes |
| N616 | Watanabe et al., Zhang et al. | Yes |
| N657 | Shajahan et al., Zhang et al., Watanabe et al. | Yes |
| N709 | Shajahan et al., Watanabe et al. | Yes |
| N717 | Shajahan et al., Watanabe et al. | Yes |
| N801 | Shajahan et al., Watanabe et al. | Yes |
| N1074 | Shajahan et al., Watanabe et al. | Yes |
| N1098 | Shajahan et al., Watanabe et al. | Yes |
| N1134 | Shajahan et al., Watanabe et al. | Yes |
| N1158 | Watanabe et al. | Yes |
| N1173 | Watanabe et al. | Yes |
| N1194 | Shajahan et al., Watanabe et al. | Yes |
| O323 | Shajahan et al. | No |
| O325 | Shajahan et al. | No |

